# An asymmetry in the frequency and position of mitosis in the epiblast precedes gastrulation and suggests a role for mitotic rounding in cell delamination during primitive streak epithelial-mesenchymal transition

**DOI:** 10.1101/2020.02.21.959080

**Authors:** Evangéline Despin-Guitard, Navrita Mathiah, Matthew Stower, Wallis Nahaboo, Elif Sema Eski, Sumeet Pal Singh, Shankar Srinivas, Isabelle Migeotte

## Abstract

The epiblast, a pseudostratified epithelium, is the precursor for the three main germ layers required for body shape and organogenesis: ectoderm, mesoderm, and endoderm. At gastrulation, a subpopulation of epiblast cells constitutes a transient posteriorly located structure called the primitive streak, where cells that undergo epithelial-mesenchymal transition make up the mesoderm and endoderm lineages.

In order to observe the behavior of individual cells, epiblast cells were labeled ubiquitously or in a mosaic fashion using fluorescent membrane reporters. The cell shapes of individual cells and the packing and behaviour of neighbouring cells during primitive streak formation were recorded through live time-lapse imaging. Posterior epiblast displayed a higher frequency of rosettes, a signature of cell rearrangements, prior to primitive streak initiation. A third of rosettes were associated with a central cell undergoing mitosis. Interestingly, cells at the primitive streak, in particular delaminating cells, underwent mitosis twice more frequently than other epiblast cells, suggesting a role for cell division in epithelial-mesenchymal transition. Pseudostratified epithelia are characterized by interkinetic nuclear migration, where mitosis occurs at the apical side of the epithelium. However, we found that exclusively on the posterior side of the epiblast, mitosis was not restricted to the apical side. Non-apical mitosis was apparent as early as E5.75, just after the establishment of the anterior-posterior axis, and prior to initiation of epithelial-mesenchymal transition. Non-apical mitosis was associated with primitive streak morphogenesis, as it occurred specifically in the streak even when ectopically located. Most non-apical mitosis resulted in one or two daughter cells leaving the epiblast layer to become mesoderm. Furthermore, in contrast to what has been described in other pseudostratified epithelia such as neuroepithelium, the majority of cells dividing apically detached completely from the basal pole in the epiblast.

Cell rearrangement associated with mitotic cell rounding in the posterior epiblast during gastrulation, in particular when it occurs on the basal side, might thus facilitate cell ingression through the PS and transition to a mesenchymal phenotype.

**GRAPHICAL ABSTRACT:** 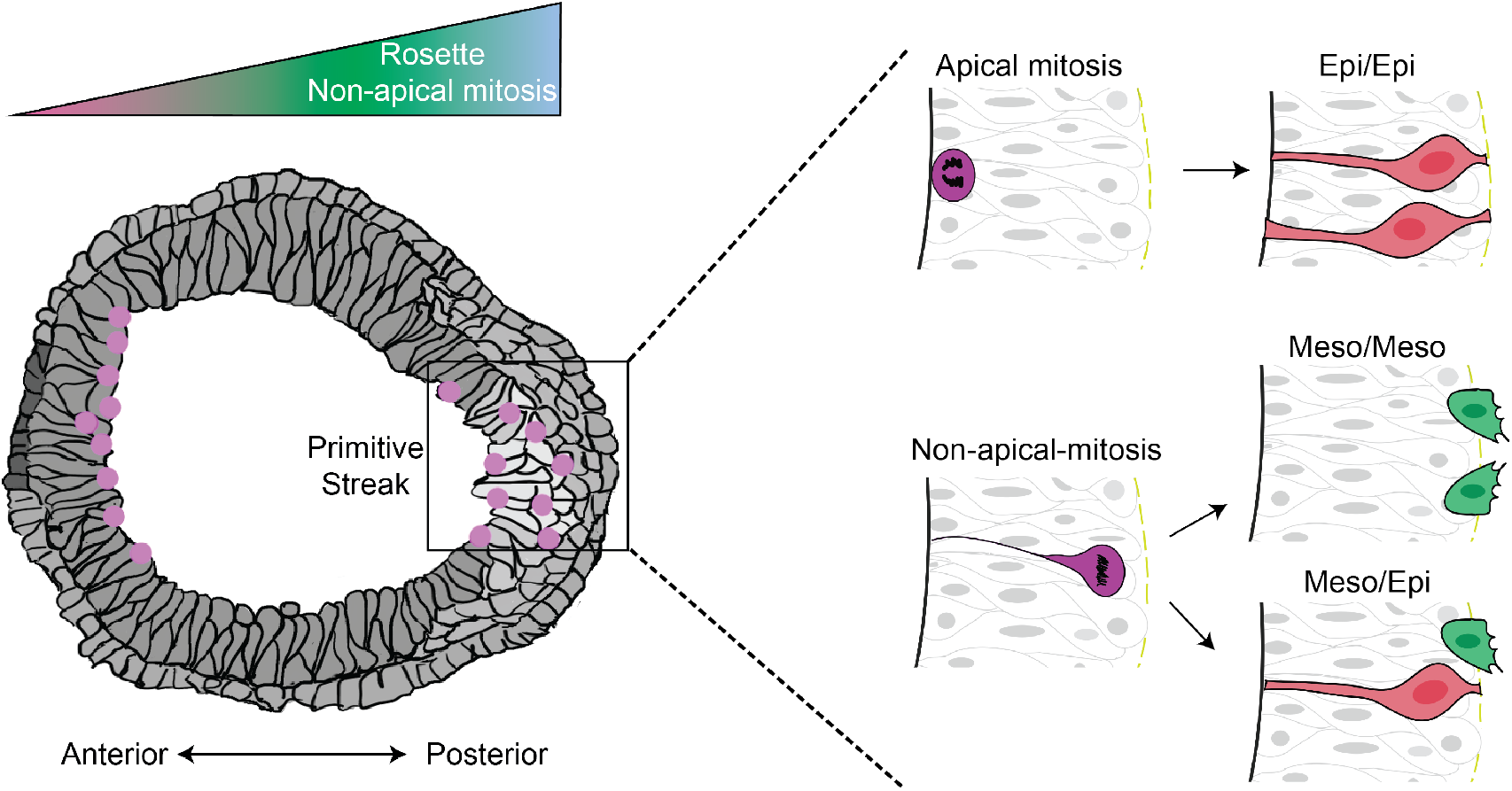

## INTRODUCTION

Gastrulation is an essential morphogenetic event that establishes the three layers of the animal body plan (ectoderm, mesoderm, and endoderm) through an epithelial-mesenchymal transition (EMT) that occurs at an organizing center called the primitive streak (PS) in amniotes (S. J. Arnold & Robertson, 2009; Ramkumar & Anderson, 2011). In mouse, the combination of multiple signaling gradients coming from embryonic epiblast, as well as extraembryonic ectoderm and visceral endoderm, creates a permissive zone in the posterior epiblast for PS establishment. The first marker of anterior-posterior asymmetry is expression of *Wnt3* in a tiny region of VE at the embryonic-extraembryonic border at embryonic day (E) 5.5 (Rivera-Pérez & Magnuson, 2005). Shortly after, between E5.5 and E5.75, a group of visceral endoderm cells at the distal end of the embryo migrates towards the embryonic-extraembryonic border in the direction opposite to *Wnt3* expression. This organizer, called the Anterior Visceral Endoderm, secretes inhibitors of Nodal and Wnt signaling that consolidate anterior-posterior axis specification (Stower & Srinivas, 2014). PS initiates in the proximal posterior epiblast at the embryonic-extraembryonic boundary, and extends distally towards the tip of the embryo. Contrary to other species, there seems to be no global epiblast movement towards the posterior side in mouse, but rather a local change of fate (Rivera-Pérez & Magnuson, 2005; Williams, Burdsal, Periasamy, Lewandoski, & Sutherland, 2012). The streak can be identified through expression of markers in pre streak (*Wnt3*), and early streak (*Eomesodermin*, and *Brachyury*) embryos at around E6 (Rivera-Perez 2005). It can be identified through morphology alone from the mid streak stage (E6.25-E6.5).

EMT allows epithelial cells to become motile through loss of cell-cell adhesion and change of polarity from apical-basal to front-rear, via cytoskeleton reorganization and acquisition of migratory and matrix remodelling capacities (Nieto, Huang, Jackson, & Thiery, 2016). Live imaging of mid and late streak embryos showed that epiblast cells in the PS region delaminate through apical constriction (leading to the so-called “bottle shape”) followed by retraction of the apical process and extrusion on the basal side (Ramkumar et al., 2016; Williams et al., 2012). Analyses of mouse mutant lines displaying an accumulation of cells within the primitive streak converge on a major role for actin-myosin cytoskeleton regulation to allow delamination at the PS (Lee, Silva-Gagliardi, Tepass, McGlade, & Anderson, 2007; Ramkumar et al., 2015, 2016). Control of cell ingression is defined at the cellular level by the complementary pattern of expression of Crumbs2 (in delaminating cells) and Myosin IIB (in their neighbors) on the apical side of epiblast cells (Ramkumar et al., 2016). *Crumbs2* mutant cells fail to detach from their basal attachment, suggesting cellular anisotropy might help extrusion from the epithelium. Interestingly, in *Crumbs2* mutant embryos, initiation of gastrulation occurs normally, and only a proportion of cells fail to ingress, leading to progressive expansion of the PS, while the rest of post-EMT cells join the mesodermal wings, suggesting that several mechanisms may control PS cellular extrusion.

An alternative delamination mechanism based on asymmetric division has been described in chick neural crest EMT from the neuroepithelium (Ahlstrom & Erickson, 2009). Similar to neuroepithelium, epiblast is a pseudostratified epithelium, and live imaging experiments have shown apical-basal nuclear movement, suggesting it undergoes interkinetic nuclear migration (IKNM) (Ichikawa et al., 2013). Classically, in IKNM, cells retain apical and basal connections, while the nucleus migrates along their apical-basal length according to cell cycle stage, and division occurs on the apical side (P J Strzyz, Matejcic, & Norden, 2016).

In order to define the cell shape changes and cellular rearrangements taking place at the prospective PS and explore alternative delamination mechanisms as gastrulation proceeds, we performed confocal and lightsheet imaging of mouse embryos expressing fluorescent reporters ubiquitously or mosaically. We found that posterior epiblast cells form rosettes with higher frequency than anterior or lateral epiblast. Dynamic and static observation of dividing cells revealed a higher frequency of mitosis in the primitive streak, and non-apical mitosis were found specifically on the posterior side of the epiblast, which resulted in ingression of one or two mesoderm cells per event.

## RESULTS

### Posterior epiblast displays higher frequency of rosettes

Lightsheet imaging, in which a thin bi-concave sheet of excitation laser optically sections an imaged sample that can be rotated and imaged at successive angles, has emerged as the method of choice for embryo live imaging, as it provides high time and space resolution tridimensional images with minimal phototoxicity (Amat & Keller, 2013). Recently, several culture methods have been developed to allow growing mouse embryos within a lightsheet microscope (Ichikawa et al., 2013; McDole et al., 2018; Udan, Piazza, Hsu, Hadjantonakis, & Dickinson, 2014). In order to record epiblast dynamics prior to, and during PS establishment, we performed lightsheet imaging at E5.75 (stage at which the anterior-posterior axis is defined) of CAG-TAG embryos bearing a membrane Tomato reporter and a nuclear (H2B) GFP reporter (Trichas, Begbie, & Srinivas, 2008), as well as *Hex*-GFP, a cytoplasmic reporter expressed in the Anterior Visceral Endoderm (to aid anterior-posterior orientation) (Srinivas, Rodriguez, Clements, Smith, & Beddington, 2004) (Fig. 1a and a’, Sup. Fig. 1a-c, and Sup. Fig. 2a). Embryos were mounted in a glass tube containing a hollow agarose cylinder. Samples were rotated in order to obtain high quality images from 4 sides, and the corresponding Z-stacks were fused to obtain a complete 3D representation. A striking feature was the abundance of rosettes, transient epithelial multicellular structures in which five or more cells interface at a central point, in the epiblast layer (Fig. 1a, Video 1). Rosettes were quantified systematically on the anterior, lateral and posterior sides on Z sections located 5 μM from the epiblast basal side. Rosette frequency was calculated through normalization according to the surface of the region of interest (in which clear cell contours could be delineated) of each section as well as total time of acquisition. We found that rosettes were twice more frequent on the posterior side (Fig. 1a and a” and Sup. Fig. 1).

**Figure 1:**
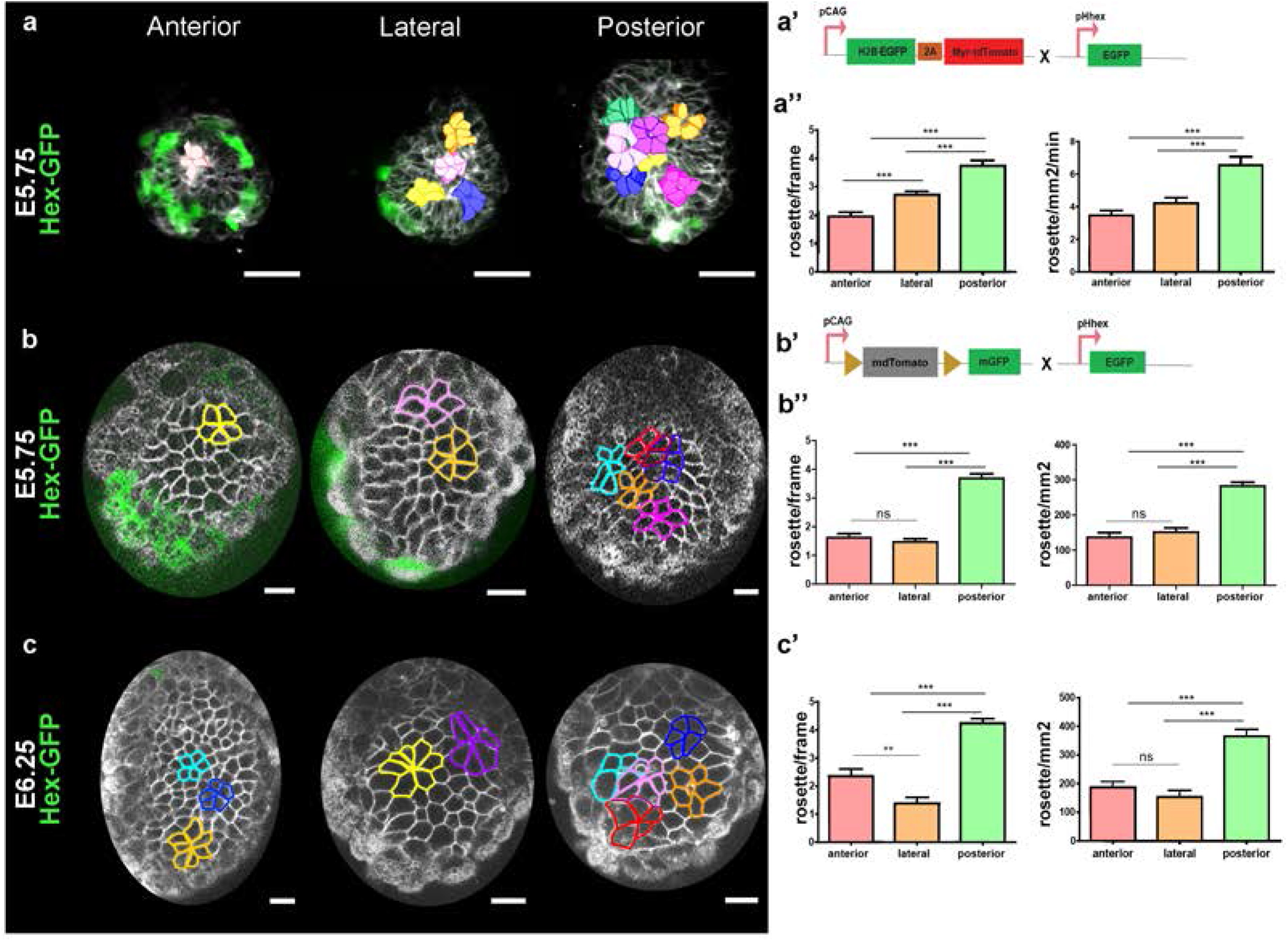
Epiblast cells organize as rosettes at the onset of primitive streak morphogenesis. (a) Manually segmented rosettes on 3D reconstructions from Z-stacks of 25 μm of an E5.75 embryo imaged using lightsheet microscopy. AVE cells are in green, and all cell membranes are labelled in grey. Left: anterior view; Middle: lateral view (anterior to the left); Right: posterior view. Scale bars: 20 μm. (a’) Mouse line strategy used to perform live imaging. (a”) Left: Quantification of the number of rosettes observed per frame. Right: Frequency of rosettes, normalized by the surface of the epiblast region in focus and the time of observation. Mean ± SEM, n=3.(b, c) Rosettes outlines on optical slices from E5.75 (b) and E6.25 (c) embryos imaged by confocal microscopy. AVE cells are in green, and all cell membranes are labelled in grey. Left: anterior view; Middle: lateral view (anterior to the left); Right: posterior view. Scale bars: 20 μm. (b’) Mouse lines strategy used to perform live imaging. (bȝ, c’) Left: Quantification of the number of rosettes observed per frame. Right: Rosettes density normalized by the surface of the epiblast region in focus. The posterior region includes the PS region, as it could not be precisely discriminated. Mean ± SEM. (b”) E5.75: n anterior=47 frames from 4 embryos, n lateral=77 frames from 9 embryos, n posterior=67 frames from 5 embryos. (c’) E6.25: n anterior=20 frames from 2 embryos, n lateral=16 frames from 1 embryo and n posterior=69 frames from 7 embryos.

The high time (7 minutes intervals) and space (1 μm Z slice) resolution allowed reconstruction of rosettes along the entire height of the epithelium, as well as follow up of individual cells fate. Around 30% of rosettes had a central cell with apical rounding, usually preceding cell division (Sup. Fig. 2b and c). However, not all apical round cells were associated with a rosette, and the frequency of apical rounding was similar on all sides (Sup. Fig. 1). Lifespan of rosettes associated with apical rounding and mitosis was longer than that of rosettes without rounding (Sup. Fig. 2c).

In order to increase sample number and to cover a longer period of development through concomitant imaging of several embryos, we performed live confocal imaging of embryos dissected at E5.75 or E6.25 (when the primitive streak is specified and cells start delaminating), and placed in optically compatible conical wells with either the anterior, posterior or lateral side facing the objective. Cell membranes were labeled ubiquitously using Rosa26::membrane dtTomato/membrane GFP (mTmG) (Muzumdar, Tasic, Miyamichi, Li, & Luo, 2007), and the *Hex*-GFP transgene was used for orientation (Fig.1b’ and Sup. Fig. 2a’). Z-stacks at 3 μm interval encompassing roughly half of the embryo were recorded with time intervals of 25 minutes. Epiblast cell contours were segmented, and rosettes were quantified as described above (Fig. 1b and c, Video 2). Due to the longer intervals between time points, normalization was performed according to surface alone. Confocal live imaging confirmed a higher frequency of rosettes on the posterior, compared to anterior and lateral epiblast (Fig. 1b” and c’), at E5.75 and E6.25.

**Figure 2:**
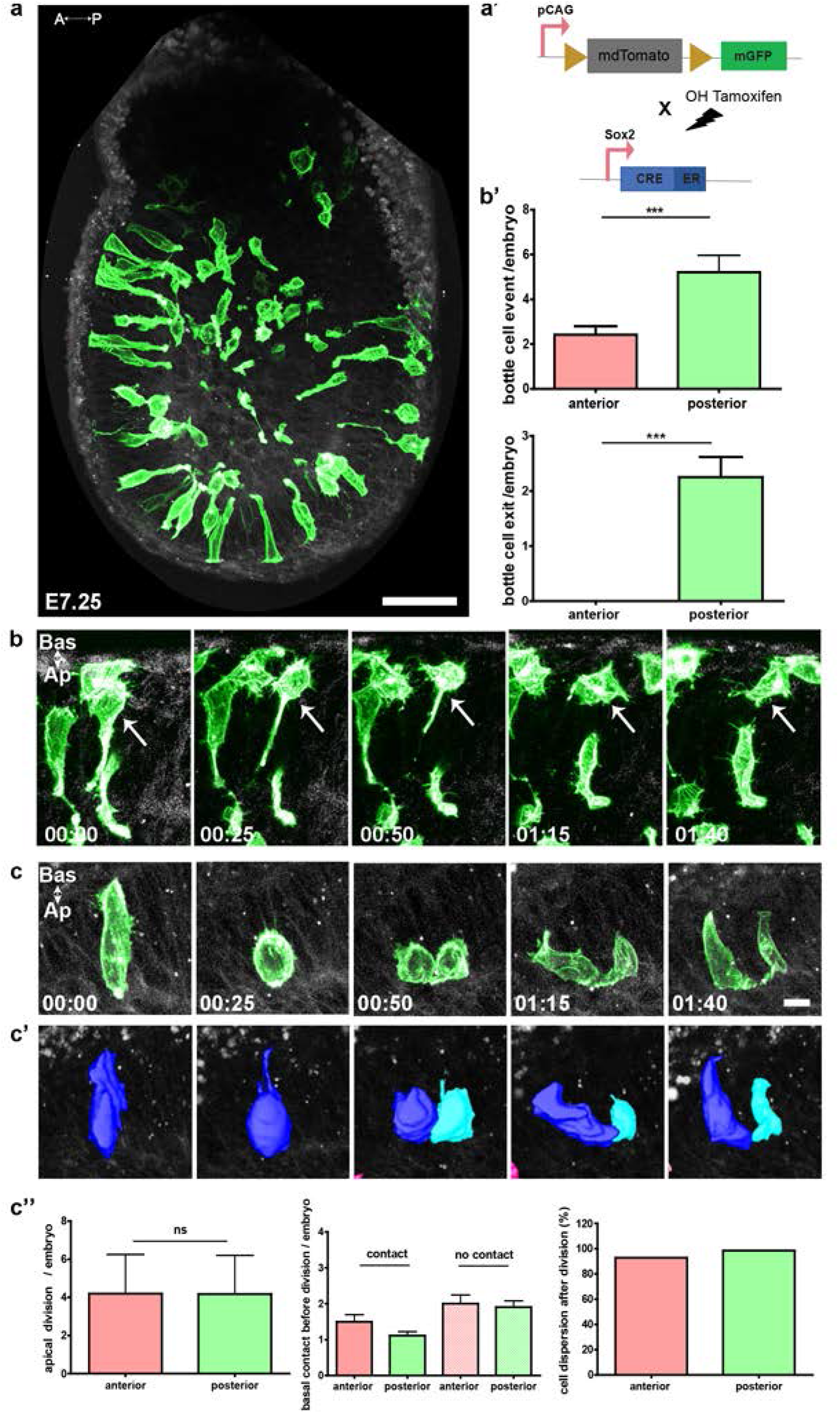
Mosaic epiblast labelling allows tracking of bottle-shaped cells and cell division. (a) Z-projection of an E7.25 embryo mosaically labelled through OH-tamoxifen injection and imaged by two-photon microscopy. Scale bar: 50 μm. n=63 embryos. (a’) Mouse line strategy used to produce embryos mosaic for membrane dtTomato (grey) and membrane GFP (green) in the epiblast. (b) Z-projections of stills from live imaging recording of a bottle-shaped cell that delaminates through the basal membrane. Scale bar: 50 μm. (b’) Quantification of bottle shape and delamination events. Mean ± SEM, n=54. Z-projection (c) and 3D reconstruction (c’) of stills from live imaging recording showing an apical division without basal contact, followed by daughter cells dispersion. Scale bar: 20 μm. (c”) Quantification of apical divisions with or without persistence of a basal contact, and their outcome in term of daughter cells dispersion. The posterior region includes the PS region, as it could not be precisely discriminated. Mean ± SEM when applicable, n=52.

The higher frequency of rosettes on the posterior side prior to the onset of gastrulation is compatible with a dynamic epithelium primed for epithelial-mesenchymal transition.

### Primitive streak cells use multiple mechanisms to delaminate

We hypothesized that the excess of rosettes on the posterior side might arise due to cell shape changes related to EMT. However, the pseudostratified and packed arrangement of the epiblast made it quite challenging to follow individual cells shape changes. OH-tamoxifen-induced recombination of mTmG through *Sox2*Cre-ERT2 (K. Arnold et al., 2013) activation allowed mosaic labeling of the epiblast (Fig. 2a, a’, Videos 3-6). At that stage of development, anterior-posterior axis can be determined through morphology alone. Due to toxicity of tamoxifen prior to implantation, we could not observe the early events of gastrulation. However, when the injection was given at E6.25, litter size and embryo morphology were normal. Live imaging was performed at E7.25, when PS elongation is complete, which offered a higher probability to observe numerous delamination events within the observation window. Embryos were placed on the lateral side in conical wells, to examine anterior and posterior epiblast. Two-photon imaging allowed recording the whole width of the embryo, with Z-intervals of 3 μm and time intervals of 25 minutes, which permitted 3D reconstruction over time. Cells acquiring either a round or a bottle shape were segmented and tracked to identify their destiny (within the epiblast or mesoderm layers) as well as that of their progeny when division occurred.

So-called “bottle-shaped” cells, with a round basal cell body and an apical extension, were present on both sides, but more frequent on the posterior side, and cell delamination only took place on the posterior side. Cells maintained an apical attachment until their basally located cell body had begun crossing the basal membrane, and only fully detached after delamination (Fig. 2b, b’).

In most cells dividing apically, a connection to the basal membrane could not be identified (Fig. 2c-c”), even when performing imaging at higher spatial resolution and laser intensity. Although we cannot exclude the possibility of a very thin cell process missed by the imaging technique, those observations are compatible with a profound reorganization in which dividing cells detached basally upon apical mitotic rounding. Daughter cells then elongated and reinserted separately in the epithelium, resulting in cell dispersion (as described in (Abe, Kutsuna, Kiyonari, Furuta, & Fujimori, 2018)). Those findings suggest that although epiblast is a pseudostratified epithelium, and undergoes IKNM, cells shape and attachment dynamics during mitosis are distinct to what is known in neuroepithelia.

For systematic quantification, epiblast regions were defined as anterior or posterior by tracing a line joining the distal pole to the middle of the embryo at the embryonic/extraembryonic border, and cells undergoing rounding were followed overtime (Fig. 3a-c). Although the frequency of cell division was similar in anterior and posterior epiblast, there was a trend towards a higher division rate specifically in cells undergoing delamination (Fig. 3d). Surprisingly, and exclusively on the posterior side, cell rounding and division happened all along the apical-basal length of the cell, so that non-apical division represented 30 % of all divisions in posterior epiblast (Fig. 3e, f). It was preferentially associated with EMT, as it resulted in formation of one or two mesoderm cells (Fig. 3g), suggesting mitotic rounding could be an alternative mechanism for cell delamination.

**Figure 3:**
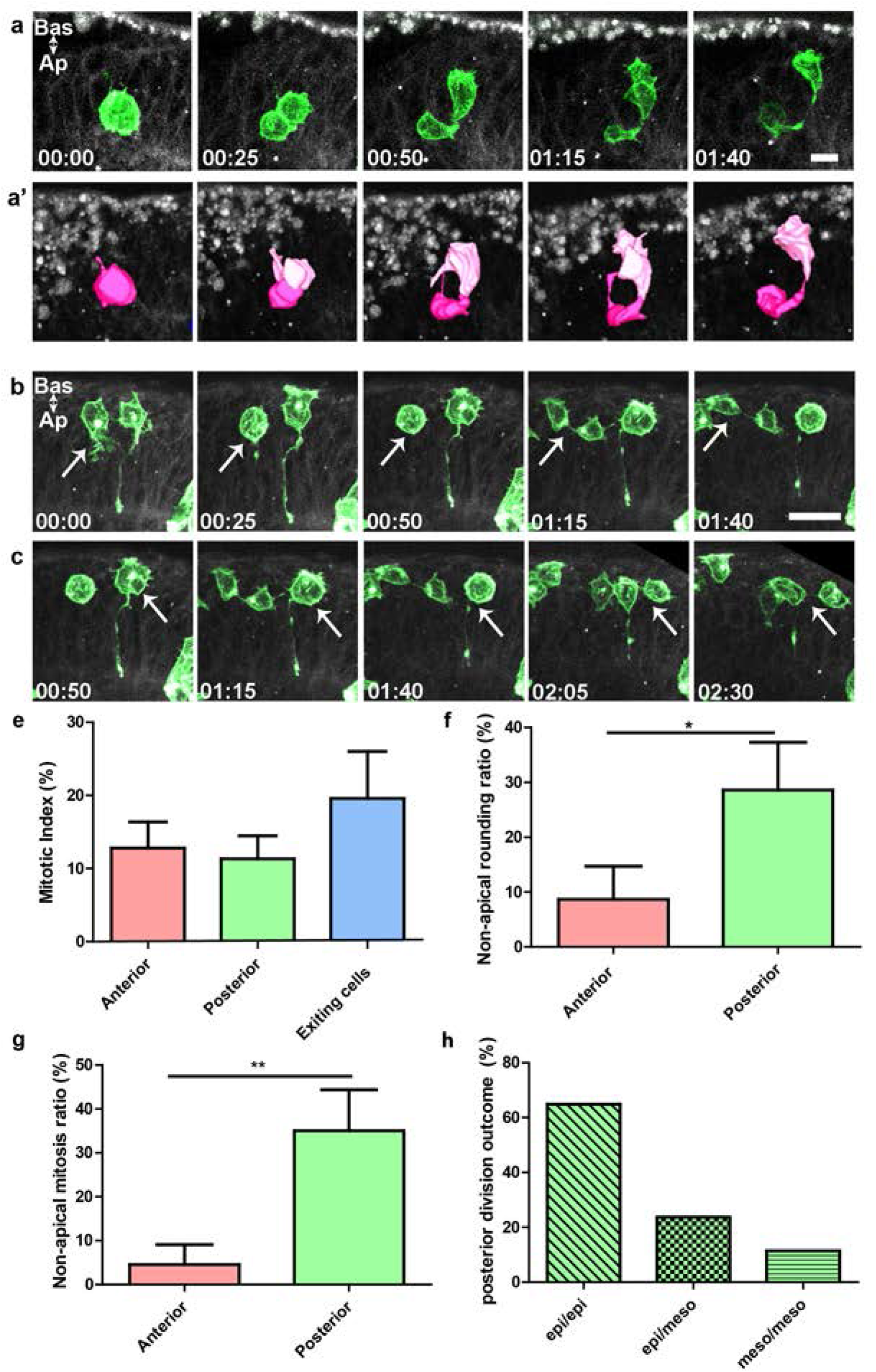
Non-apical division occurs in the posterior epiblast and can give rise to one or two mesoderm cells. Z-projection (a) and 3D cell reconstruction (a’) of stills from two-photon live imaging of an E7.25 embryo mosaically labelled for epiblast cells showing a division giving rise to an epiblast and a mesoderm cell that exits from the epiblast layer. Scale bar represents 20 μm. (b) (c) Z-projection of stills from two-photon live imaging of an E7.25 embryo mosaically labelled for epiblast cells. Non-apical division can be observed in the posterior epiblast only, without (b) or with (c) attachment to the apical side. Scale bar: 20 μm (d) Mitotic index (green dividing cells/ total number of green cells) in the anterior and posterior region, as well as among cells exiting the epiblast at the primitive streak. The posterior region includes the PS region, as it could not be precisely discriminated. (e) Ratio of non-apical rounding (non-apical rounding/ total rounding) normalized by the number of green cells and expressed in percentage for the anterior and posterior regions. (f) Ratio of non-apical division (non-apical division/ total division) normalized by the number of green cells and expressed in percentage for the anterior and posterior region of the embryo. (g) Quantification of posterior cell divisions outcomes: either 2 epiblast daughter cells (epi/epi), one epiblast and one mesoderm daughter cells (epi/meso) or 2 mesoderm daughter cells (meso/meso). Data are normalized by the number of green cells. n=43 embryos.

### Higher frequency of mitosis, including non-apical mitosis, in primitive streak and pre-gastrulation posterior epiblast

In order to document the proportion and position of mitotic nuclei all along the gastrulation period, we performed immunostaining for phospho-histone H3 on sections or whole-mount embryos from E5.75 to E7 using lightsheet and confocal imaging (Fig. 4 and 5). Samples were counterstained for nuclei (DAPI) for cell count and F-actin (Phalloidin) for cell shape, and we quantified mitotic index, apical-basal position of positive nuclei, as well as, when possible, orientation of mitotic plate relative to apical-basal axis.

**Figure 4:**
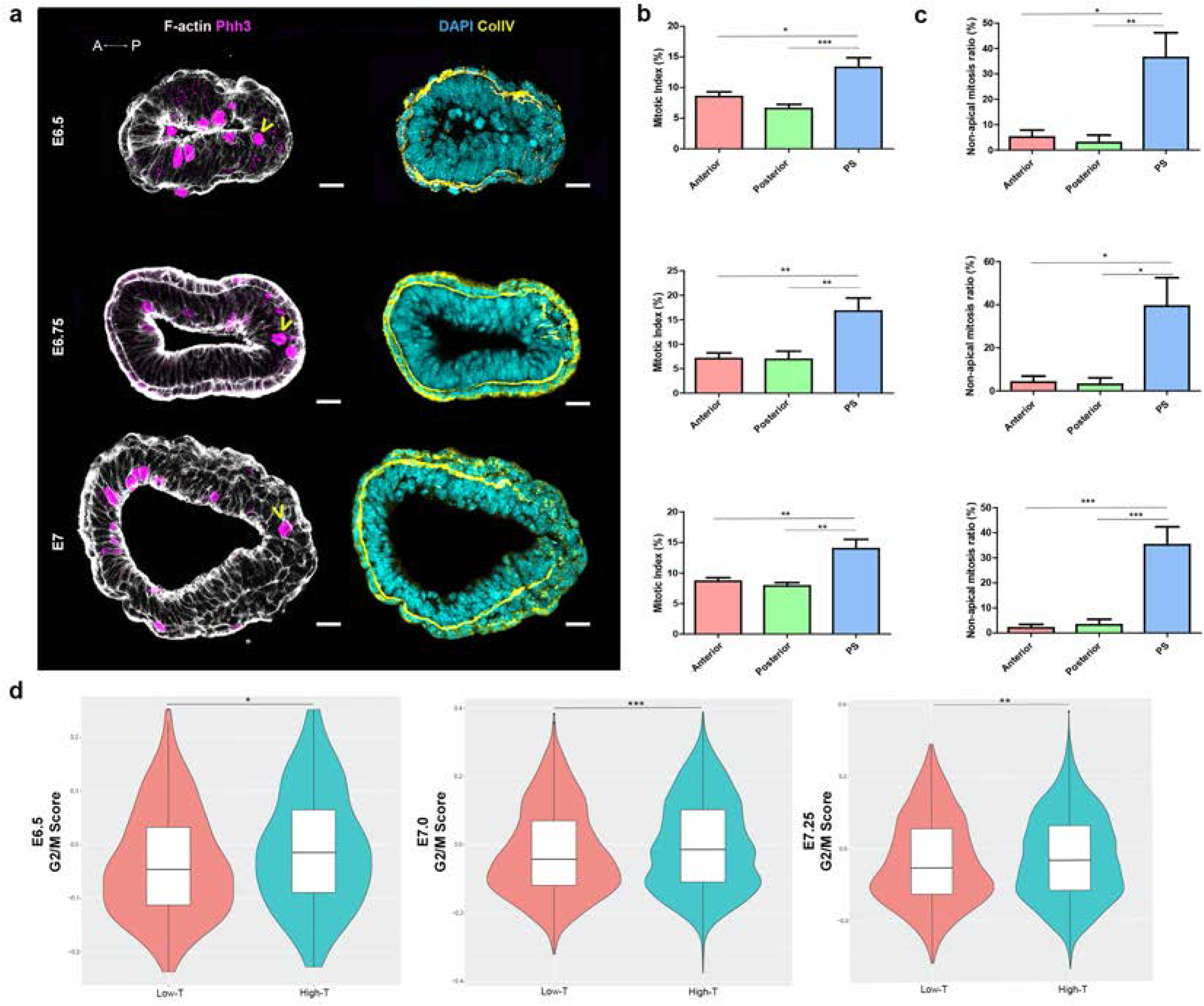
Mitosis is more frequent at the primitive streak, and primitive streak mitotic rounding is not always apical. (a) Z-projections of transversal sections from E6.5 (top), E6.75 (mid) and E7 (bottom) embryos. Samples were stained for F-actin (Phalloidin, grey) and mitosis (Phh3, magenta) on the left and nuclei (DAPI, cyan) and basal membrane (collagen IV, yellow) on the right. Yellow arrowheads show non-apical mitosis. Scale bar: 25 μm. (b) Mitotic index (Phh3/DAPI) in percentage at the anterior, posterior, and primitive streak (PS) regions of E6.5 (top), E6.75 (mid) and E7 (bottom) embryos. (c) Ratio of non-apical mitosis (non-apical Phh3/ Phh3, normalized by number of nuclei) in percentage at the anterior, posterior, and PS regions of E6.5 (top), E6.75 (mid) and E7 (bottom) embryos. The PS region is defined by the area where the basal membrane (yellow) is degraded, and the posterior region quantification excludes counts from the PS region. Values are shown as Mean ± SEM. E6.5: n=20 frames from 6 embryos, E6.75: n=13 frames from 5 embryos, E7: n=20 frames from 6 embryos. (d) Violin plots representing the distribution of cells in G2/M phase in cells with high, compared to low, *Brachyury* expression among cells annotated as primitive streak and mesoderm harvested from E6.5, E7, and E7.25 embryos. A: anterior, P: posterior, PS: primitive streak

**Figure 5:**
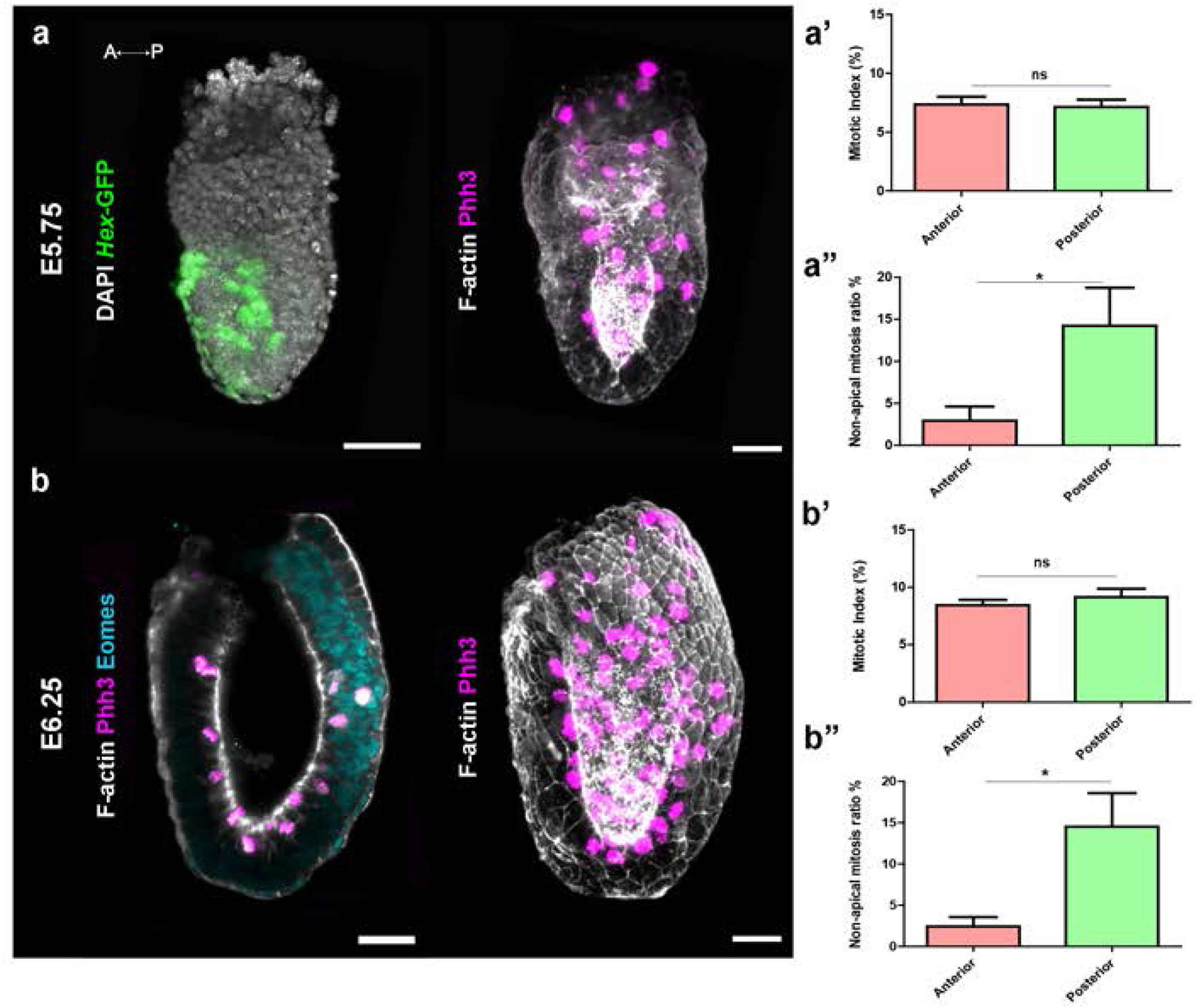
Non-apical divisions occur in the posterior epiblast before and at the onset of gastrulation. (a) 3D reconstruction of wholemount lightsheet imaging of an E5.75 *Hex*-GFP (green, anterior marker) embryo, stained for nuclei (DAPI, grey) on the left, mitosis (Phh3, magenta) and F-actin (Phalloidin, grey) on the right. (b) Z-slice (left) and 3D reconstruction (right) of wholemount lightsheet imaging of an E6.25 embryo stained for a posterior marker (Eomesodermin, cyan), mitosis (Phh3, magenta) and F-actin (Phalloidin, grey). Scale bar: 50 μm. (a’, b’) Mitotic index (Phh3/DAPI) in percentage at the anterior and posterior regions of E5.75 (a’) and (b’) E6.25 embryos. (a”,b”) Ratio of non-apical mitosis (non-apical Phh3/Phh3, normalized by number of nuclei) in percentage in the anterior and posterior region of E5.75 (a”) and E6.25 (b”) embryos. Posterior region includes the PS region, as it could not be precisely discriminated. Values are shown as Mean ± SEM. E5.75: n=11 embryos, E6.75: n=6 embryos.

We first focused on early to mid streak embryos (E6.5, E6.75, and E7). Based on anatomy and localization of Collagen IV, a basal membrane marker, we defined anterior (anterior half), posterior (posterior half except PS area), and PS region (where basal membrane is perforated) on transverse sections (Fig. 4a and Sup. Fig. 3a). Remarkably, mitosis was around 2 times more frequent in the PS region compared to the rest of the epiblast (Fig. 4b). In addition, over a third of mitotic nuclei were located away from the apical surface in PS cells (Fig. 4c). Contrary to apical mitosis, the majority of phospho-histone H3 positive cells on the basal side retained a connection with the apical side (Sup. Fig. 3b). We found no preferential mitotic plate orientation (Sup. Fig. 3c). At the PS, the mitotic index and the proportion of non-apical mitosis were stable overtime (Sup. Fig. 3d-f). The higher frequency of mitosis in the streak was corroborated by examining signatures of cell cycle phases in single cell transcriptomes from the Mouse Gastrulation 2018 atlas (Pijuan-Sala et al., 2019). We selected cells annotated as PS and mesoderm at E6.5, E7, and E7.25. At each time point, cells with high *Brachyury* expression, which likely represent cells at the PS since *Brachyury* is progressively down-regulated in nascent mesoderm, had a higher probability to display a G2/M signature (Fig.4d and Sup. Fig. 4).

For earlier embryos, anterior-posterior orientation was determined either through *Hex*-GFP (Anterior Visceral Endoderm, Fig. 5a) or immunostaining for Eomesodermin (PS, Fig. 5b), and embryos were examined in whole-mount in a sagittal orientation. The proportion of non-apical mitosis was significantly higher on the posterior side, even before the primitive streak could be identified morphologically (Fig. 5a’, a”, b’, b”).

### Non-apical mitosis is associated with primitive streak morphogenesis

PS specification occurs at the region of the embryo opposite the Anterior Visceral Endoderm: when the Anterior Visceral Endoderm remains distal, PS forms a ring at the embryonic/extraembryonic boundary, and when it migrates partially, PS is skewed to the posterior side but fails to elongate (Rakeman & Anderson, 2006). To assess whether non-apical mitosis is a feature systematically associated with the PS, we examined embryos displaying defective Anterior Visceral Endoderm migration resulting in an ectopic primitive streak. E6.25 *Rac1*^*KO*^ (Migeotte, Omelchenko, Hall, & Anderson, 2010) or *RhoA*^*VE-deleted*^ (Christodoulou et al., in press) embryos were immunostained for Cerberus1 (an Anterior Visceral Endoderm marker), phospho-histone H3, and counterstained for nuclei (DAPI) and F-actin (Phalloidin). In both models, Anterior Visceral Endoderm migration was impaired (Fig. 6a-d and Sup. Fig. 5a-d). Non-apical mitosis was significantly and consistently more frequent in the region of the embryo most distant from the Anterior Visceral Endoderm (Fig. 6b, c, e and Sup. Fig. 5b, c, e), showing it is specifically associated with PS morphogenesis, rather then with a geographic position in the embryo.

**Figure 6:**
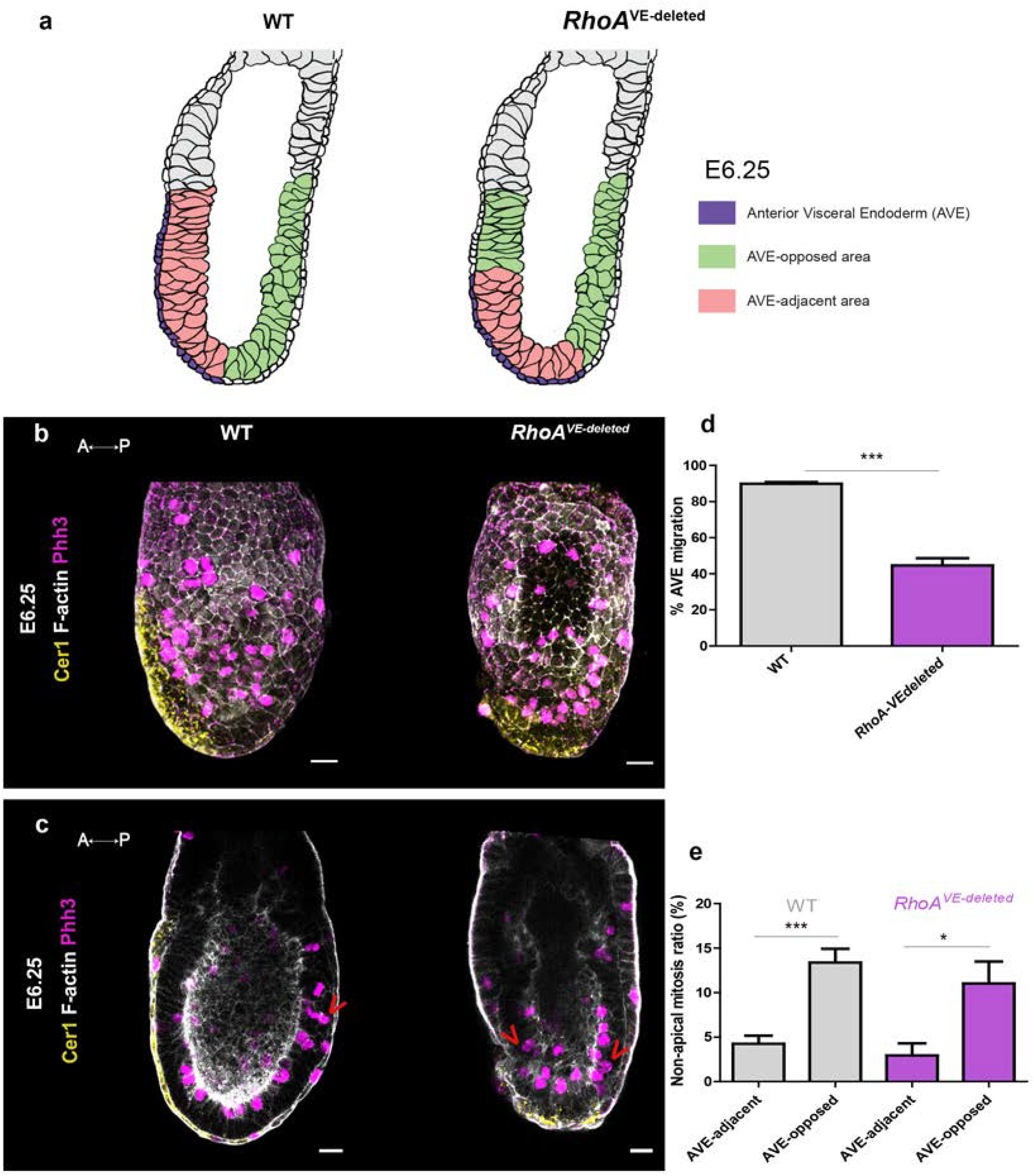
Non-apical mitosis is a feature associated with the primitive streak. (a) Representation of the position of the AVE-opposed area (green) and the AVE-adjacent area (pink) in WT (left) and mutants (right) embryos. (b, c) 3D reconstruction (b) and confocal Z-slice (c) of E6.25 WT (left) and *RhoA*^VE-deleted^ (right) embryos stained for a AVE marker (Cerberus 1, yellow), F-actin (Phalloidin, grey) and mitosis (Phh3, magenta). Scale bar: 25 μm. Red arrowheads point to non-apical mitosis. (d) Percentage of AVE migration in WT and *RhoA*^VE-deleted^ mutants. (e) Ratio of non-apical mitosis (non-apical Phh3/ Phh3, normalised by the volume in μm^3^) in percentage, in AVE-adjacent and AVE-opposed regions from WT and *RhoA*^VE-deleted^ embryos. Values are shown as Mean ± SEM. WT: n=31; *RhoA*^VE-deleted^: n=17.

## DISCUSSION

Anterior-Posterior axis specification of the mouse embryo precedes detection of PS markers by half a day, and initiation of EMT by a day. Through lightsheet and confocal live imaging of embryos that had completed Anterior Visceral Endoderm migration, we observed a high frequency of rosettes in the posterior epiblast, prior to and during PS morphogenesis. Rosettes are intermediate stages of epithelial reorganization observed in multiple morphogenetic events in diverse organisms (Harding, McGraw, & Nechiporuk, 2014). There are two main classes of rosettes, characterized by their mechanisms of formation and resolution. Rosettes arising through planar polarized constriction are usually short lived, and often contribute to tissue elongation. On the other hand, rosettes arising through apical constriction persist for longer periods of time, and do not resolve in a stereotyped fashion but rather remodel to participate to the formation of a structure or organ, often through generation of a fold or lumen. Nodal-dependent rosettes with actin-rich centers have been observed within developing PS of the chick embryo (Wagstaff, Bellett, Mogensen, & Münsterberg, 2008), and they are more frequent as development progresses (Yanagawa, Sakabe, Sakata, Yamagishi, & Nakajima, 2011). However, their mode of formation is unclear. In view of the complex pseudostratified organization of the epiblast, we could not identify rosettes mode of formation with precision. A proportion of rosettes related to collapse of the basal process during apical mitotic rounding resulting in bridging of the surrounding cells. In later stages, rosettes were likely formed in a similar fashion (although with an inverse apical-basal polarity) through apical constriction of delaminating bottle-shaped cells. However, since we observed rosettes prior to initiation of EMT, it is likely that a proportion of those arise through a yet to discover mechanism, possibly linked to the need for increased epithelium fluidity during PS morphogenesis. The recent advances of mouse embryo lightsheet imaging (McDole et al., 2018), including dynamic 3D reconstruction of multicolor embryos, will likely allow to resolve the different modes of rosette formation in the PS along gastrulation.

Through mosaic labeling of the epiblast, we followed cell shape changes in mid to late streak embryos. Cells delaminating through apical constriction retained an apical connection until their cell body had exited the epiblast layer. Interestingly, cells leaving the PS appeared to have a higher frequency of division. This was confirmed by a systematic quantification of mitotic nuclei in anterior, posterior, and primitive streak epiblast, showing higher mitotic index in cells adjacent to the perforated segment of the basement membrane. Data mining from single cell transcriptomes (Pijuan-Sala et al., 2019) highlighted a higher probability to be in G2/M phase for PS and mesoderm cells expressing high level of Brachyury. Similarly, it has been shown in rat embryos that cells in the PS region cycled more rapidly than the rest of the epiblast (Mac Auley, Werb, & Mirkes, 1993). Together, those observations point towards a role for mitosis in EMT at the PS. A potential mechanistic explanation comes from recent work in *Drosophila* showing that pseudostratified epithelial cells down-regulate E-cadherin as they round up for mitosis (Aguilar-Aragon, Bonello, Bell, Fletcher, & Thompson, 2020).

In addition, we found difference in localization of mitotic rounding and cell division depending on anterior-posterior localization in the epiblast. In anterior epiblast, mitotic rounding was universally apical. Different to other pseudostratified epithelia, in the majority of dividing cells we could not distinguish a basal attachment, suggesting cells detach from the basal membrane prior to division. In posterior epiblast, and particularly in the streak area, mitotic rounding was not exclusively apical, but could occur all along apical-basal cell length. In mutant embryos where primitive streak was ectopic due to Anterior Visceral Endoderm migration failure, non-apical division was detectable in the region of the epiblast further away from the Anterior Visceral Endoderm, hence associated to PS morphogenesis. Remarkably, non-apical division resulted in extrusion of one or two daughter cells. Non-apical mitosis has been described in chick dorsal neural tube, where neural crest delamination occurs (Ahlstrom & Erickson, 2009): live imaging showed that a proportion of rounded mitotic cells detached from the lumen to undergo mitosis, and that daughter cells from those non-apical mitosis all became neural crest. In the zebrafish retina, perturbation of CDK1-driven apical IKNM resulted in non-apical mitosis and disrupted integrity of the pseudostratified neuroepithelium because of aberrant cell delamination (Paulina J. Strzyz et al., 2015). In Drosophila tracheal placode, mitotic rounding is necessary to accelerate epithelial invagination (Kondo & Hayashi, 2013). Similarly, cell division is critical for chick embryo gastrulation via its role in epithelial rearrangements through regulation of cortical actomyosin (Firmino, Rocancourt, Saadaoui, Moreau, & Gros, 2016).

In light of those results, we propose that increase in epithelial fluidity, redistribution of cell contacts through acquisition of the round morphology associated with mitosis, as well as, possibly, mechanical pressure on the basal membrane from non-apical dividing cells, could facilitate transition to a mesenchymal phenotype at the PS.

## ACKNOWLEDGMENTS

We thank the animal house and light microscopy (LiMiF) core facilities at the ULB (Erasme Campus). We thank M. Martens and J-M. Vanderwinden for help with confocal imaging. N.M. and E.D.G. received a FRIA fellowship of the Fonds de la Recherche Scientifique (FNRS), N.M. was also supported by the “Fonds David et Alice van Buuren” and the “Fondation Jaumotte-Demoulin”. W.N. was supported by WELBIO (SGR2015). S.S. is funded through Wellcome Senior Investigator Award 105031/C/14/Z. S.P.S. is supported by the FNRS under grant number 34772792 (MISU). I.M. is a FNRS research associate. WELBIO, the FNRS, and the Fondation Erasme supported this work. The authors declare no financial or non-financial competing interests.

## METHODS

### Mouse strains and genotyping

The mTmG (Muzumdar et al., 2007) and the *Sox2*-Cre-ERT2 (K. Arnold et al., 2013) lines were obtained from the Jackson Laboratory, the CAG-TAG (Trichas et al., 2008) and the *Hex*-GFP (Srinivas et al., 2004) lines from Shankar Srinivas. Mice were kept on a CD1 background. Mice colonies were maintained in a certified animal facility in accordance with European guidelines. The local ethics committee (CEBEA) approved all experiments. OH-tamoxifen (Sigma) was suspended at 100 mg/ml in ethanol 100%, and diluted in sesame oil (Sigma) to a final concentration of 10 mg/ml. Females were injected intraperitoneally with 2 mg OH-tamoxifen at E6.25 and E6.75. Embryos were collected at E7.25.

Mouse genomic DNA was isolated from ear biopsies following overnight digestion at 55°C with 1.5% Proteinase K (Quiagen) diluted in Lysis reagent (DirectPCR, Viagen), followed by heat inactivation.

### Embryo culture and live imaging

#### Confocal Imaging

Embryos were dissected in Dulbecco’s modified Eagles medium (DMEM) F-12 (Gibco) supplemented with 10% FBS and 1% Penicillin-Streptomycin and L-glutamine and 15 mM HEPES. They were then cultured in 50% DMEM-F12 with L-glutamine without phenol red, 50% rat serum (Janvier), at 37°C and 5% CO_2_. Embryos were observed in suspension in individual conical wells (Ibidi) to limit drift, under a Zeiss LSM 780 microscope equipped with Plan Apochromat 25x/0.8, C Achroplan 32x/0.85, and LD C Apochromat 40x/1.1 objectives. Stacks were acquired every 25 minutes with 3 μM Z intervals for up to 10 hours. Embryos were cultured for an additional 6 to 12 hours after imaging to check for fitness.

#### Lightsheet Imaging

20 μL glass capillaries (Brand, 701904) with plungers were used to create a 2% agarose cylinder, in which a copper wire of 150 μm (for E5.75 embryo) or 195 μm (for E6.5 embryos) was inserted. Once agarose was solidified, the wire was removed to leave a tunnel in the agarose cylinder, in which the embryo was placed.

Embryos were dissected in dissection medium, and allowed to recover in equilibrated culture medium (50% CMRL Medium, 50% KO Serum, and 0,02% glutamine) in an incubator (37°C, 5% CO2) for 1 h. Embryos were transferred to a Petri dish filled with culture medium in which the agarose cylinders were lying flat, and were gently moved into the hollow cylinder. Once the cylinders were mounted into glass capillaries, those were placed into a syringe adapted with tips on both ends to secure the capillary.

The lightsheet microscope (Zeiss Lightsheet Z.1) chamber was filled with culture medium, and left to equilibrate with the sample holder at 37°C and 5% CO2 prior to imaging for 1h. Imaging was performed using a 20x, NA1.0 Plan-Apochromat water immersion objective, and dual side illumination. For CAG-TAG embryos, 1 to 2% of laser power (488 nm, 543 nm) was applied with an exposure time of 29 ms. Embryos were imaged on 4 sides (anterior, posterior, 2 laterals) with an interval of 90°. Stack Images were taken in dual illumination. Images were captured at a 1024×1024 resolution, with Z intervals of 1 μm for maximum 200 μm, and time intervals of 7 minutes. Embryos were cultured for up to 12 hours.

### Antibodies

Antibodies were: rat anti-Ph3 (Abcam; 1/500), rabbit anti-Tbr2/Eomes (Abcam; 1/100), goat anti-Brachyury (R&D Systems; 1/20), Goat anti-collagenIV (Millipore, 1/500), Goat anti-Cerberus1 (R&D, 1/500), Rabbit anti-Phh3 (Sigma, 1/500). F-actin was visualized using 1.5 U/ml TRITC-Phalloidin (Invitrogen; 1/100), and nuclei using DAPI (Sigma; 1/1000). Secondary antibodies were: anti goat Alexa Fluor 647, anti-goat Alexa Fluor 488, anti-rabbit Alexa Fluor 488, anti-rat Alexa Fluor 543, anti-rat Alexa Fluor 647, all at 1/500 (Jackson).

### Embryo Analysis

For immunofluorescence, embryos were fixed in PBS containing 4% paraformaldehyde (PFA) for 2 hours at 4°C, cryopreserved in 30% sucrose, embedded in OCT and cryosectioned at 7-10 μM. Staining was performed in PBS containing 0.5% Triton X-100, 0,1% BSA and 5% heat-inactivated horse serum. Sections and whole-mount embryos were imaged on a Zeiss LSM 780 or Lightsheet Z.1 microscope.

### Image analysis

Lightsheet Z-stacks from 4 sides were fused using Zeiss plugin for Lightsheet Imaging. Images were then processed using Arivis Vision4D v2.12.3 (Arivis, Germany). Embryo contours were segmented manually on each Z-slice and time point. Embryo rotation was adjusted manually if necessary. For 3D reconstruction, cells were manually segmented on each Z-slice and time point by highlighting cellular membranes using Wacom’s Cintiq 13HD. For quantification, rosettes were manually annotated and counted on Z sections located 5 μm from the basal side of the epiblast. Presence of apical rounding on the corresponding apical side was assessed for each rosette. For Phospho-histone H3 quantifications, sections were chosen at least 10 μm apart to ensure that each cell was only counted once, and counting was performed using the Icy software (http://icy.bioimageanalysis.org). Videos were generated using the Arivis Vision4D software.

For each population, normality was assessed using a Shapiro-Wilk test. According to the results of the precedent test, samples were compared using a non-parametric Mann-Whitney test or an unpaired t-test. Ns: non-significant, *: P-value≤0.05, **: P-value≤0.01 and ***: P-value≤0.001.

### Single-cell RNA-seq Analysis

To mine the Mouse Gastrulation Atlas (Pijuan-Sala et al., 2019), the scRNA-seq data was downloaded using the ‘MouseGastrulationData’ package (Griffiths & Lun, 2019). The data include raw gene counts and annotations for each cell (cell-type, stage and UMAP-coordinates). To plot the data, we utilized the UMAP-coordinates, cell-type and stage annotations provided by the authors. We subset the data by selecting cells annotated as primitive streak, nascent mesoderm, extraembryonic mesoderm, mixed mesoderm and intermediate mesoderm from E6.5, E7.0 and E7.25 stages. Raw data was normalized for library size and mitochondrial counts and scaled using the ‘SCTransform’ function from Seurat 3.1 (Hafemeister & Satija, 2019). Cell cycle score was calculated from scaled data using ‘CellCycleScoring’ function in Seurat package (Butler, Hoffman, Smibert, Papalexi, & Satija, 2018). This function predicts the cell phase for each cell using G2/M and S phase markers provided in the package, and assigns each cell a quantitative G2/M and S score. These scores and predicted phase of each cell are stored in Seurat object meta data. Further, after assigning cells as high-T or low-T based on the expression level of *Brachyury* (cut-off on scaled expression: >0.5), UMAP and violin plots of G2M scores were plotted in R. Statistical analysis was performed in R (Student’s t-test).

**VIDEOS LEGENDS**

**Supplementary Video 1: Rosette formation in epiblast.** Z sections from lighsheet microscopy live imaging of a E5.75 embryo expressing *Hex*-GFP (green) and membrane Tomato (grey) acquired from the anterior (left), lateral (middle) and posterior (right) sides. Rosettes are highlighted in color. Interval time: 7 min and scale bar: 25 μm.

**Supplementary Video 2: Rosette dynamics in posterior epiblast.** Z sections from confocal microscopy live imaging of a E5.75 embryo expressing *Hex*-GFP (green) and membrane Tomato (grey) placed on its posterior side. Rosettes are highlighted in color. Interval time: 20 min and scale bar: 50 μm.

**Supplementary Videos 3-6: Cell shape and cell division tracking in mosaically labelled epiblast.** 3D reconstructions from two-photon microscopy live imaging of mTmG; *Sox2*Cre-ERT2 E7.25 embryos where epiblast cells are mosaically labelled through OH-tamoxifen injection at E6.25. Interval time: 20 min. Scale is indicated by a grid in μm at the beginning of the videos.

---

**Supplementary Figure 1:**
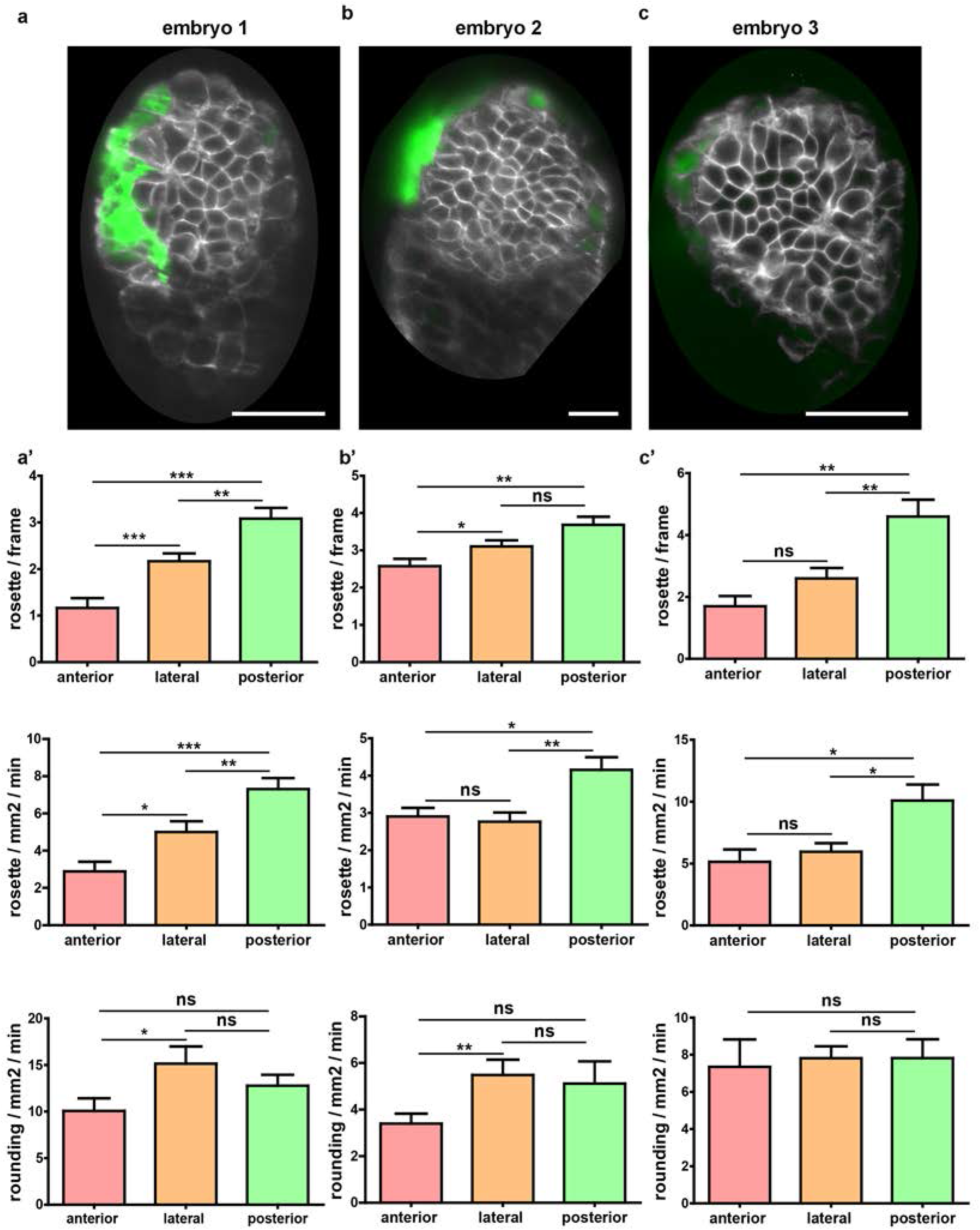
Quantification of rosette events in three pre-gastrulation embryos using lightsheet imaging. (a, b, c) Single optical slices from 3 embryos at E5.75. All embryos are in lateral orientation, with the anterior side to the left as defined by the position of AVE cells (Hex-GFP, in green). Membranes are marked by dtTomato (in grey) (a’) (b’) (c’) Upper row: Quantification of the number of rosettes observed per frame; Mid row: Frequency of rosettes, normalized by the surface or the epiblast region in focus and the time of observation; Bottom row: Frequency of apical cell rounding, normalized by the surface or the epiblast region in focus and the time of observation. Data show consistently a higher number of rosettes in the posterior side compared to the anterior and lateral side, both in absolute and normalized values. Time interval between acquisition is 7 min, and interval between Z-sections is 1 μm. Values are shown as Mean ± SEM, and number of time points analysed per embryo are n(a’)=12, n(b’)=19 and n(c’)=10. Scale bars represent 20 μm in (a) and (c), and 50 μm in (b).

**Supplementary Figure 2:**
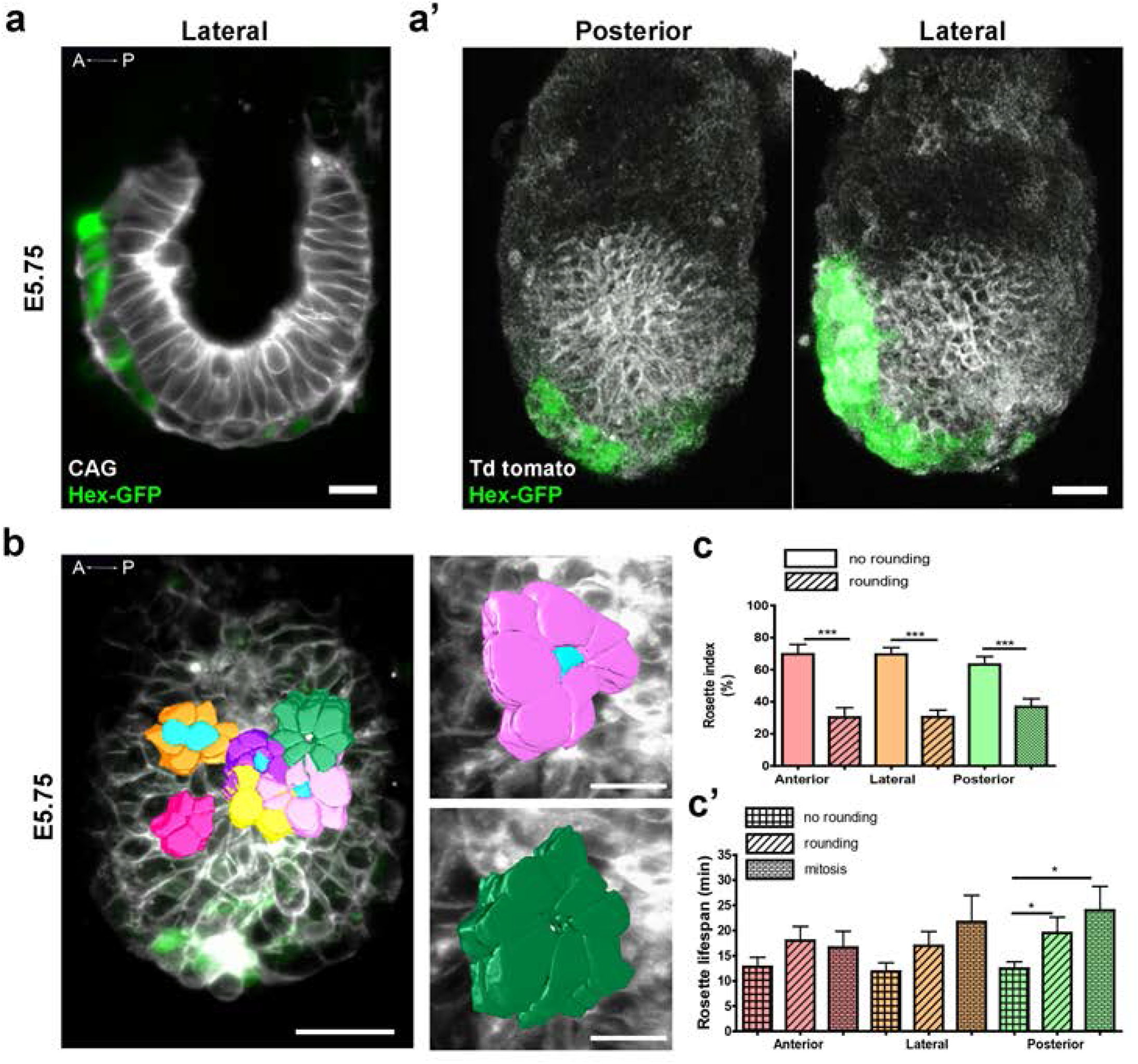
Embryo orientation for lightsheet and confocal imaging, and quantification of rosettes linked to apical cell rounding. (a) Z-projection of the middle section of an embryo imaged by lightsheet microscopy, shown in a lateral view with anterior to the left. Cell shape can be easily segmented. Scale bar: 10 μm. (a’) 3D rendering of an embryo imaged by confocal microscopy, shown in posterior (left) and lateral (right) views. Scale bar represents 50 μm. In (a) and (a’), AVE cells are identified through the Hex-GFP reporter, allowing embryo antero-posterior orientation. (b) Left panel: 3D rendering of the apical side of manually segmented rosettes from an embryo imaged using lightsheet microscopy. Scale bar: 50 μm. Highlighted in light blue are rounded cells localised at the centre of the rosette. Right panels: zooms of a rosette linked to a rounded apical cell (top) and a rosette without an apically rounded cell (bottom). Scale bar: 20 μm. (c) Rosette rounding indexes: rosettes with or without central apical rounded cell /total rosettes in percentage and (c’) Lifespan (min) of rosettes when bound to a central mitotic, central round or non-round cell in the anterior, lateral and posterior regions of the embryo. The posterior region includes the PS region. Values are shown as Mean ± SEM.

**Supplementary Figure 3:**
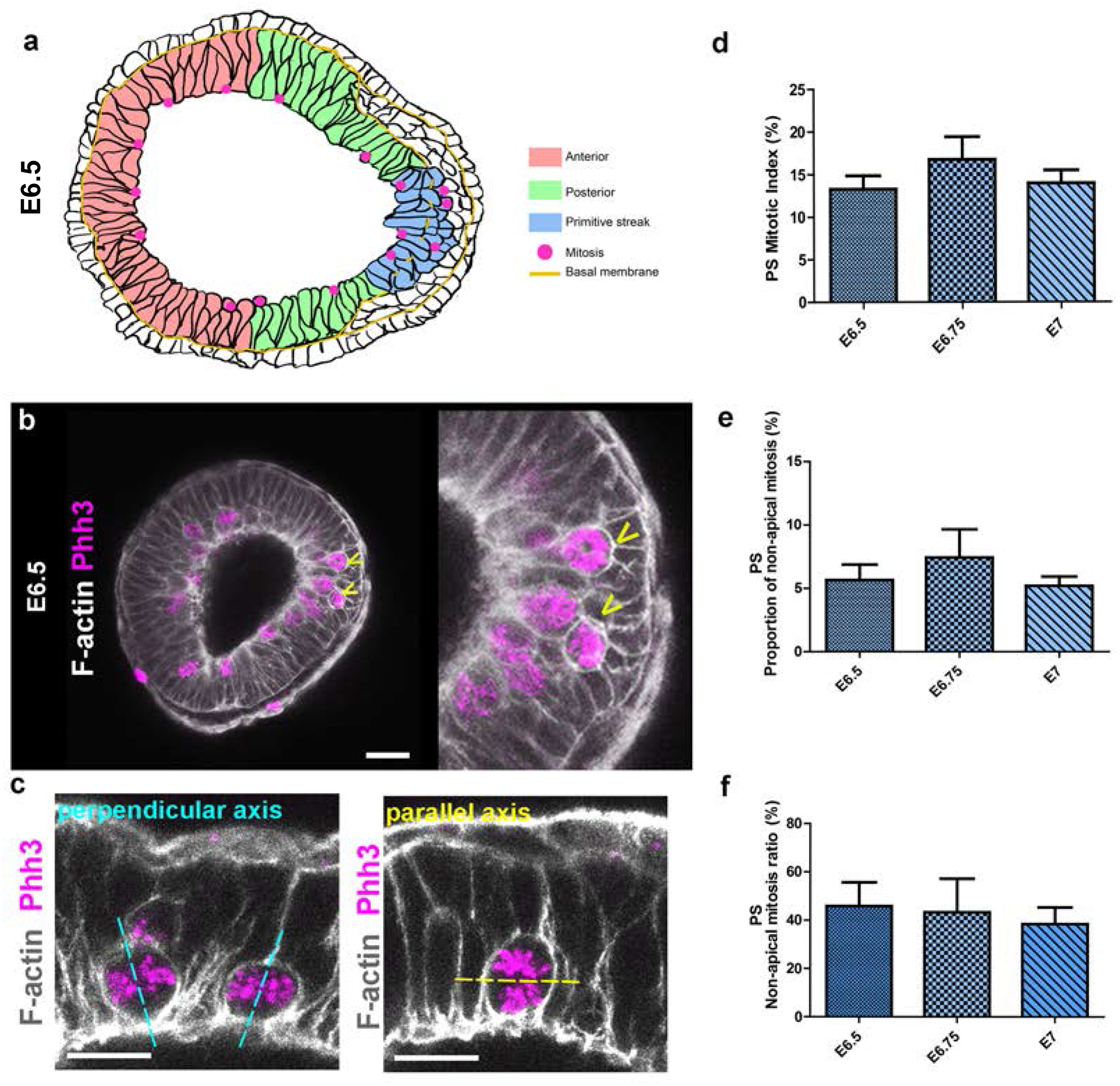
Non-apical division at the PS. (a) Representation of sagittal section from an E6.5 embryo. The epiblast is divided in three distinct regions: anterior (red), posterior (green) and primitive streak (PS, blue). The PS region is defined by the area where the basal membrane (yellow) is degraded. A mitosis is considered non-apical when occurring away from the epiblast apical border at a minimum distance corresponding to the size of one nucleus (± 10 μm), in a cell that has not crossed the perforated basal membrane and does not display a mesenchymal phenotype. (b) Transversal optical section at streak level from a E6.5 embryo labelled for mitosis (Phh3, magenta) and F-actin (Phalloidin, grey). The apical cell attachment of non-apical Phh3-positive cells is highlighted in the zoom. Scale bar: 50 μm. Arrowheads show non-apical mitoses.(c) Zoom of confocal imaging section of E6.25 embryos showing mitoses (Phh3, magenta) occurring with a parallel (left) or a perpendicular (right) axis at the apical pole. Scale bar: 30 μm. (d, e, f) Proportion of mitosis and non-apical mitosis is stable through time at the PS: Graphs representing (d) mitotic index (Phh3/ DAPI) at the PS of E6.5, E6.75 and E7 embryos. (e) Proportion of non-apical mitosis (Non-apical Phh3/ DAPI, in percentage), and (f) Ratio of non-apical mitosis (Non-apical Phh3/ Phh3 normalized by the number of nuclei, in percentage. Values are shown as Mean ± SEM. E6.5: n=20 frames from 6 embryos, E6.75: n=13 frames from 5 embryos, E7: n=20 frames from 6 embryos.

**Supplementary Figure 4:**
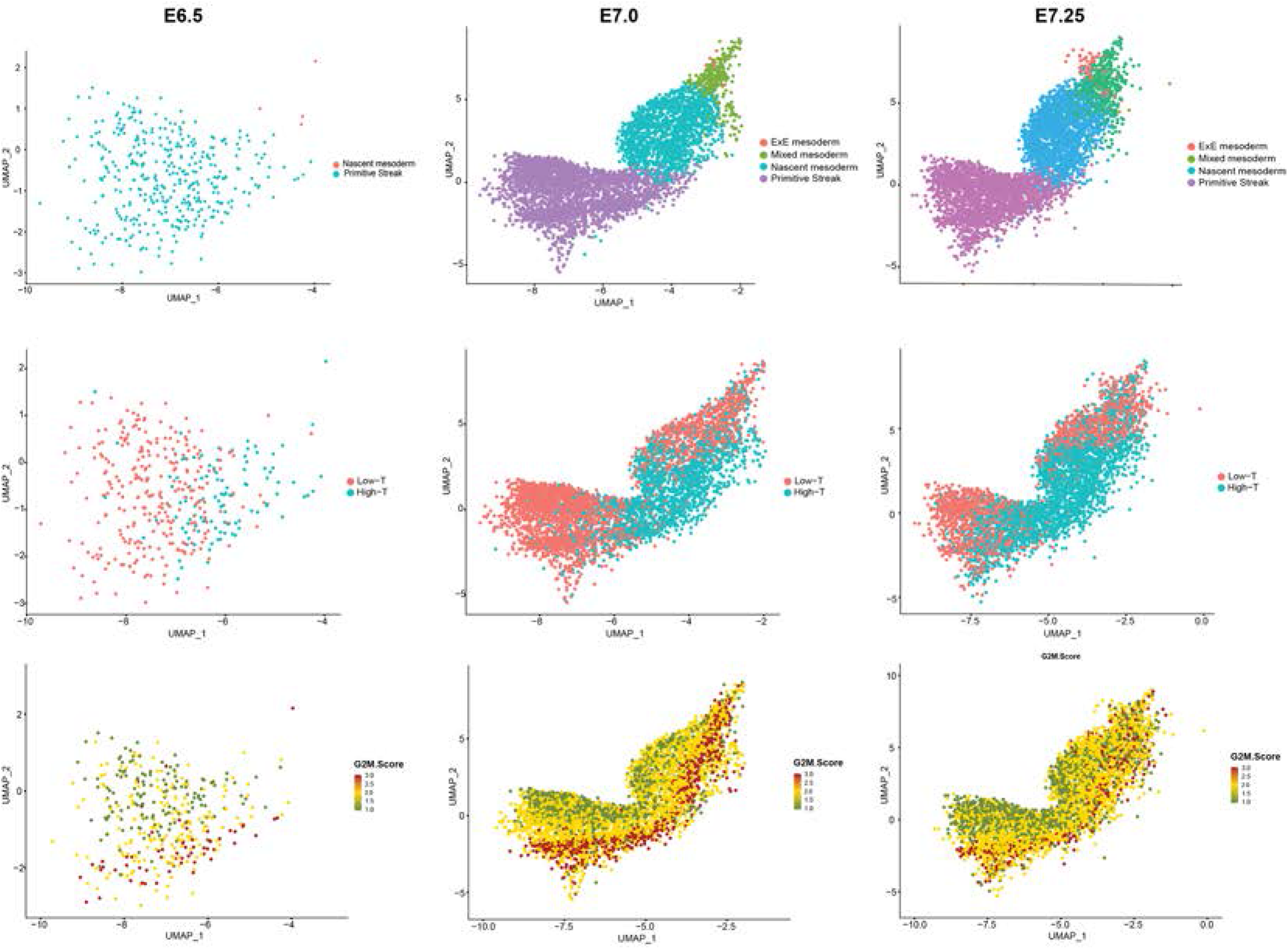
Distribution of cell cycle phases among primitive streak and mesoderm cells. UMAP plots were generated from the Mouse Gastrulation 2018 atlas. Cells annotated as primitive streak and mesoderm were selected from E6.5, E7 and E7.25 samples. Cells are coloured based on annotation in (a), on level of *Brachyury* expression in (b), and on score for G2/M signature in (c).

**Supplementary Figure 5:**
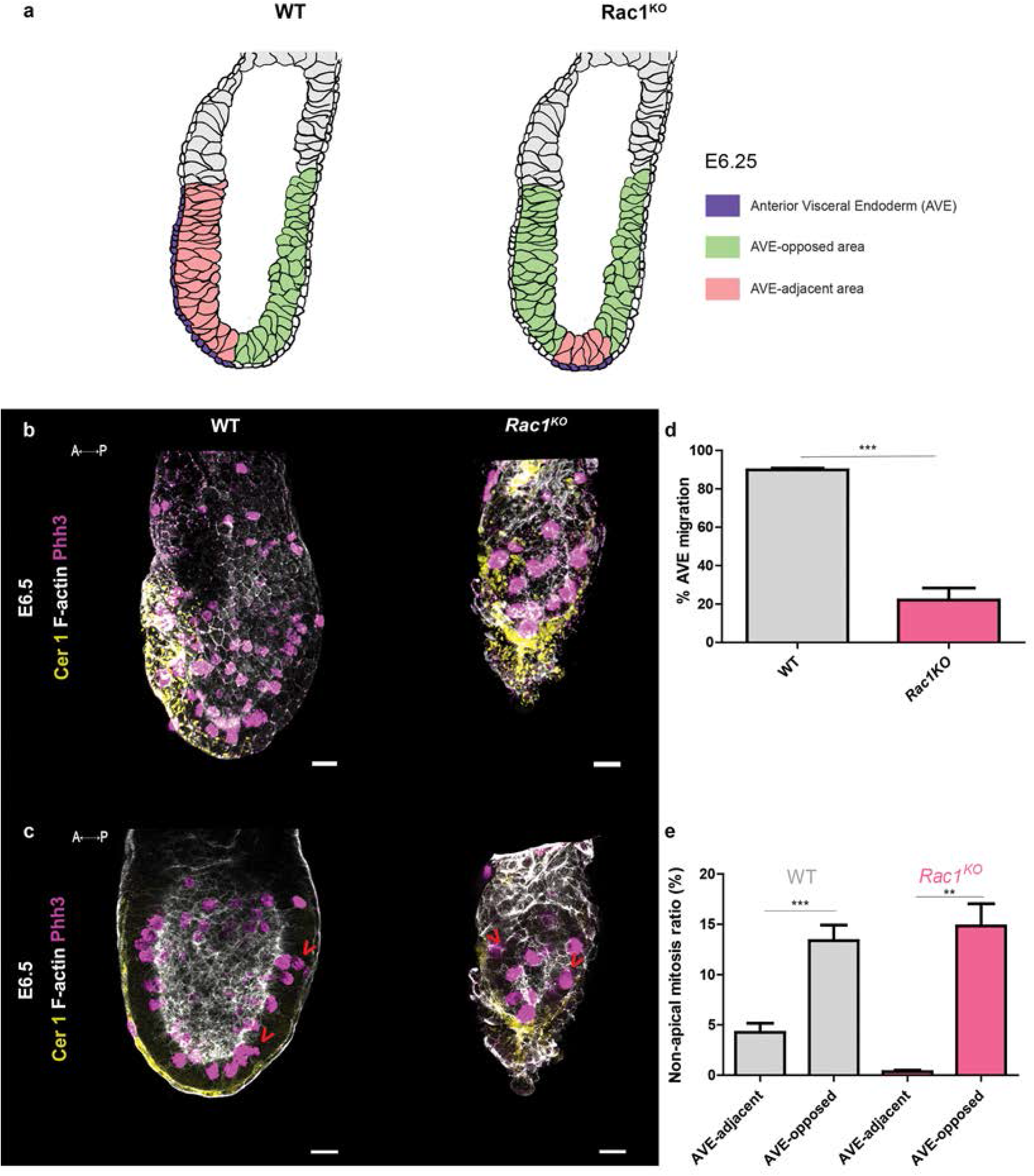
Non-apical mitosis is a feature associated with the primitive streak. (a) Representation of the position of the AVE-opposed area (green) and the AVE-adjacent area (pink) in WT (left) and mutants (right) embryos. (b, c) 3D reconstruction (b) and confocal Z-slice (c) of E6.5 WT (left) and *Rac1*^KO^ (right) embryos stained for a AVE marker (Cerberus, yellow), F-actin (Phalloidin, grey) and mitosis (Phh3, magenta). Scale bar: 25 μm. (d) Percentage of AVE migration in WT and *Rac1*^KO^ mutants. (e) Ratio of non-apical mitosis (non-apical Phh3/ Phh3 normalised by the volume) in percentage in WT and *Rac1*^KO^ mutants in AVE-adjacent compared to AVE-opposed regions. Values are shown as Mean ± SEM. WT: n=30; *Rac1*^KO^: n=7.

